# 3D Printed Customizable Radiopaque Markers for Assessing Gastrointestinal Transit

**DOI:** 10.64898/2026.05.19.726145

**Authors:** Yusheng Zhang, Parima Phowarasoontorn, Maylis Boitet, Abdel-Hameed Dabbour, Heba T Naser, Batoul Khlaifat, Khalil B Ramadi

**Affiliations:** Division of Engineering, New York University Abu Dhabi, Abu Dhabi, UAE; Tandon School of Engineering, New York University, New York, NY 11201, USA; Grossman School of Medicine and NYU Langone Health, New York University, New York, NY 11201, USA; Center for Brain and Health, New York University Abu Dhabi, Abu Dhabi, UAE; Center for Translational Medical Devices, New York University Abu Dhabi, Abu Dhabi, UAE; Core Technology Platforms, New York University Abu Dhabi, Abu Dhabi, UAE

## Abstract

Tracking gastrointestinal (GI) transit in preclinical models is essential for assessing gut motility and drug delivery. Current preclinical methods rely on end-to-end transit measurements or emptying studies that require terminal endpoints and organ explanation. Clinically, radiopaque “Sitz” markers are administered orally and their position in the GI tract is assessed through radiography. Sitz markers have been in use since 1969 and are typically mass-produced using industrial molding or extrusion, resulting in a single, fixed geometry with limited tunability. We present a stereolithography (SLA)-based method to fabricate customizable radiopaque markers using additive manufacturing with a barium sulfate (BaSO_4_)-doped resin. We demonstrate precise control over marker geometry, a key advantage over existing markers. Furthermore, we apply this method in vivo, tracking markers in a live rat model from ingestion to excretion using serial CT imaging. We systematically investigate how changes in marker geometry impact GI residency and transit time. Our results show that 3D printed markers provide a flexible and tunable platform for radiopaque marker fabrication and enable investigation of the fundamental relationship between a marker’s physical properties and its performance in a dynamic biological environment. This work establishes a novel, tunable platform for GI motility evaluation and drug delivery studies.

## 1. Introduction

Gastrointestinal (GI) motility is a fundamental physiological process that dictates the rate of digestion, nutrient absorption, and waste transit. A thorough understanding of GI motility is vital for the diagnosis and treatment of a range of GI disorders, such as gastroparesis, chronic constipation, and irritable bowel syndrome, as well as the design of oral drug delivery systems. One standard clinical approach to evaluate GI motility relies on conventional radiopaque “Sitz” markers. Patients ingest a capsule containing radiopaque markers, and their movement is tracked over time with radiography (Figure 1A). The number of markers remaining help differentiate normal, rapid, and delayed transit (Figure 1B). These markers were first developed in 1969 and have remained unchanged since, with a single geometry utilized across patients^1^. Investigation into how a marker’s physical properties, such as shape and size, affect its residency within a dynamic biological system, has been largely unexplored.

**Figure 1.**
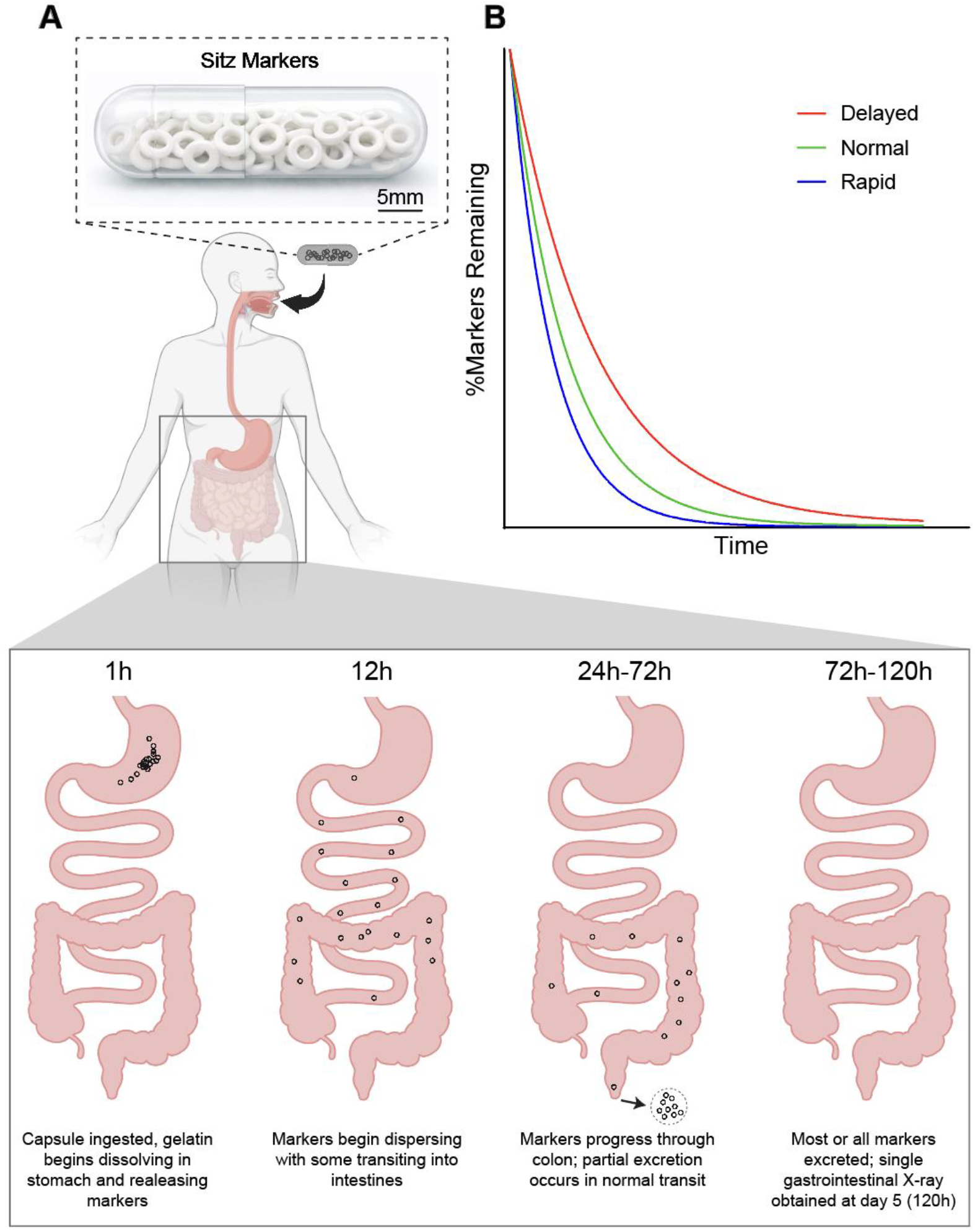
Conventional sitz marker testing and comparison to 3D-printed markers. (A) Photograph of a capsule containing multiple conventional radiopaque sitz markers.(B) Schematic representation of clinical sitz marker testing, showing ingestion of a marker-containing capsule and the use of serial imaging to distinguish rapid, normal, and delayed gastrointestinal transit based on marker clearance over time^2^.

Sitz markers were developed and validated primarily in the context of constipation and delayed colonic transit, where prolonged marker retention provides a robust diagnostic signal, and are correspondingly less informative in settings where transit is rapid and retention is minimal^4^. In such contexts, marker platforms with customizable physical properties may provide a means to better interrogate transit behavior by allowing marker design to be tuned to different physiological regimes, without altering established counting or imaging protocols. Moreover, it is also important to study how marker geometry influences gastrointestinal residency, as an improved understanding of these physical determinants could inform strategies to modulate marker retention time within the GI tract. Foundational work established that properties such as size, shape (testing rings, tetrahedrons, and clovers), and polymer flexibility are critical factors that dictate the gastric retention time of a marker, as observed via X-ray in canine models^5^. Building on these principles, modern additive manufacturing techniques enable more sophisticated designs such as microcontainers with mucoadhesive geometries^6^.

Here, we design and fabricate customizable SLA-printed barium sulfate (BaSO_4_) markers for real-time GI transit tracking in a live rat model. We systematically evaluate how the marker’s physical properties and geometry affect residency time within the GI tract. We find that systematic variation of marker geometry is associated with differences in gastrointestinal residency, accompanied by distinct deformation and stress profiles observed during transit. Together, this work establishes a novel and adaptable platform for preclinical GI motility research.

## 2. Materials and Methods

### 2.1 Marker Fabrication

We fabricated radiopaque markers using a stereolithography (SLA) 3D printer (Formlabs Form 3+). Custom photopolymer resin was prepared by incorporating barium sulfate (BaSO4, Sigma-Aldrich, USA) into a clear methacrylate-based resin (Formlabs Flex80A). Resin mixtures were prepared at a BaSO4 concentration of 35% w/w. We modified a Form 3+ 3D printer to enable more economical resin testing by reducing the minimum required resin volume. Inspired by the work of^7^, we designed a custom tank divider using SolidWorks, which we then printed on the Form 3+ with Clear V4 resin. This rectangular frame, with dimensions of 50 × 60 × 25 mm (W, L, H), was secured to the resin tank’s membrane using a silicone sealant, which was allowed to air dry for 24 hours. A custom built platform made of Aluminum was fabricated via CNC machining, with its top fixture printed on a Stratasys J750. This platform was designed to have four separate building areas, each measuring 30 x 50 mm.

Markers were designed in SolidWorks as hollow cylindrical structures. STL files were exported and printed at 0.05 mm layer thickness. After printing, the markers were washed in isopropanol to remove excess resin and cured under UV at 60 degrees C for 15 minutes submerged in water.

### 2.2 In Vivo Study

All animal experiments were conducted in accordance with protocols approved by the Institutional Animal Care and Use Committee (IACUC) at New York University (approved protocol number NYUAD-210007). Animals were housed in standard individually ventilated cages under a 12-hour light–dark cycle, with ad libitum access to food and water. Adult male Sprague Dawley rats (60 weeks old, n = 4-6, Charles River) were used in this study. Following oral administration, the animals were single housed until all markers had been recovered from the feces to ensure traceability.

### 2.3 Marker Quantification

Barium sulfate doped markers (35% BaSO_4_) were 3D printed in a control geometry (1mm OD, 0.58mm ID, 1mm length) and in three experimental variations, 2x OD/width (2xW), 5x length (5xL), or 2x OD/width and 10x length (2xW 10xL). For the in vivo study, each rat was administered an equal number of control markers and *one* of the three experimental marker types via oral gavage, enclosed in a size 9 or 9el capsule. Gastrointestinal (GI) transit was monitored using a SkyScan 1276 microCT system (Bruker, USA). Animals were anesthetized with 3-3.5% isoflurane in 0.8 L/min oxygen and positioned in dorsal decubitus on the scanner bed, secured with colored tape to minimize respiratory motion.

MicroCT imaging was performed at 75 μm spatial resolution (image resolution 512 × 1008) with X-ray source settings of 85 kV and 200 μA, using an Al+Cu filter and a 1.2° rotation step for a complete 360° rotation. Images were reconstructed with NRecon software (Bruker, USA) using no smoothing, a 20% beam hardening correction, and a ring artifact correction of 2. MicroCT scans were acquired at defined intervals: 1 hour post gavage to confirm administration, 7–9 hours to capture early GI distribution, 24 hours, and subsequently every 24 hours until all markers had been excreted. Fecal samples were collected and scanned to confirm marker excretion and to recover markers for further analysis. Retention was quantified as the percentage of markers remaining in the GI tract relative to the number initially administered.

### 2.4 Marker Quantification

Scanning electron microscopy (SEM) was used to assess marker surface morphology and structural integrity before administration and after gastrointestinal transit. Marker were rinsed with deionized water, air dried overnight, and mounted on aluminum stubs using carbon tape. Sample were sputter coated with a thin layer of gold to minimize charging. SEM imaging was performed using a Thermo Fisher Scientific Quanta 3D scanning electron microscope operated at an accelerating voltage of 10kV with a spot size of 3. Images were acquired at multiple magnifications to evaluate surface features, hollow core preservation, and evidence of cracking, swelling, or structural degradation.

### 2.5 Uniaxial Compression Testing

Uniaxial compression testing was performed using an Instron mechanical testing system equipped with a 5N load cell to characterize the mechanical response of the printed markers. Individual markers were placed horizontally between two rigid, parallel compression platens and subjected to displacement controlled compression at a constant rate of 6mm/min. Compressive force and displacement were recorded continuously throughout the test. All tests were conducted at room temperature, and multiple samples were test for each geometry (n=4 per group). The resulting compressive force was converted to an effective pressure using an estimated contact area, and effective pressure was subsequently compared as a function of prescribed displacement.

## 3. Results

### 3.1 Influence of Geometry on GI Residency

We sought to understand and model the mechanics of forces experienced by the ingested devices. We first considered a simplified physical model describing how a cylindrical object interacts with the intestinal wall during transit. The axial drag *F*_*drag*_ exerted by the gut is proportional to the wall shear stress acting on the marker’s contact surface:

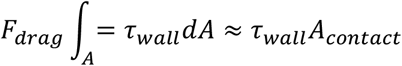

Where *τ*_*wall*_is the shear stress generated by intestinal contractions, and *A*_*contact*_is the surface area of the marker in contact with the lumen wall.

For a cylindrical marker with outer diameter *D*_*marker*_and length L, this contact area is approximately the curved surface area of the cylinder:

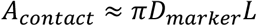

Where *πD*_*marker*_is the circumference of the marker Substituting yields:

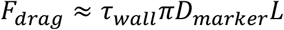

Because any swallowed object must be smaller than the intestinal inner diameter *D*_*lumen*_, we have:

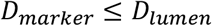

Where *D*_*lumen*_ is the physiologically determined maximum lumen diameter during normal distension.

Near occlusive markers approach this maximum, so their effective diameter is limited by the intestine’s ability to expand:

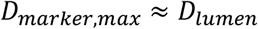

Under this physiological constraint, the diameter becomes effectively fixed in vivo, and axial drag scales primarily with the marker’s length:

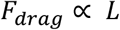

Based on this analysis, we hypothesized that marker length is a key contributor to GI residency time due to increased axial drag, where longer markers experience greater resistance to peristaltic transport along the intestinal axis, resulting in prolonged retention. This hypothesis is counter-intuitive in the context of conventional emphasis on marker width or deformability, yet it provides a mechanistic basis for understanding why elongated geometries may exhibit prolonged retention despite comparable or greater deformability.

To evaluate how each marker responds to peristaltic compression, we simulated radial squeezing of the markers using a simplified boundary condition. Each marker was placed horizontally between two opposing rigid plates, and a uniform compressive pressure was applied along its length. This setup mimics the axial component of peristaltic forces that acts to propel intraluminal contents forward. The pressure values were selected based on previously reported rat intraluminal pressure measurements with resting pressures of 2-4 mmHg and propulsive wave peaks of approximately 9±1 mmHg^8^. We applied pressures of 2,5,7, and 9mmHg (converted to Pa) to each marker design to span physiologically relevant range of intestinal peristalsis.

We began with the existing Sitz marker dimension of 1mm outer diameter, 0.58mm inner diameter, and 1mm length. We then adapted this to generate markers with 2x the width (inner diameter unchanged), 5x the length, and a combined 2x width and 10x length. Across all applied pressures, the von Mises stress distribution for the control geometry exhibited the highest stresses, with the 5xL marker showing slightly lower stresses than the control. Both width-modified geometry (2xW and 2xW 10xL) showed substantially lower stresses than the control and 5xL, with 2xW 10xL consistently exhibiting the lowest stresses (Figure 2A-H). These trends are also reflected in the peak stress-pressure curves (Figure 2J) and in the peak deformation-pressure curves (Figure 2K), where the ordering of peak von Mises stress magnitudes and deformation across the pressure range is: control > 5xL > 2xW > 2xW 10xL.

**Figure 2.**
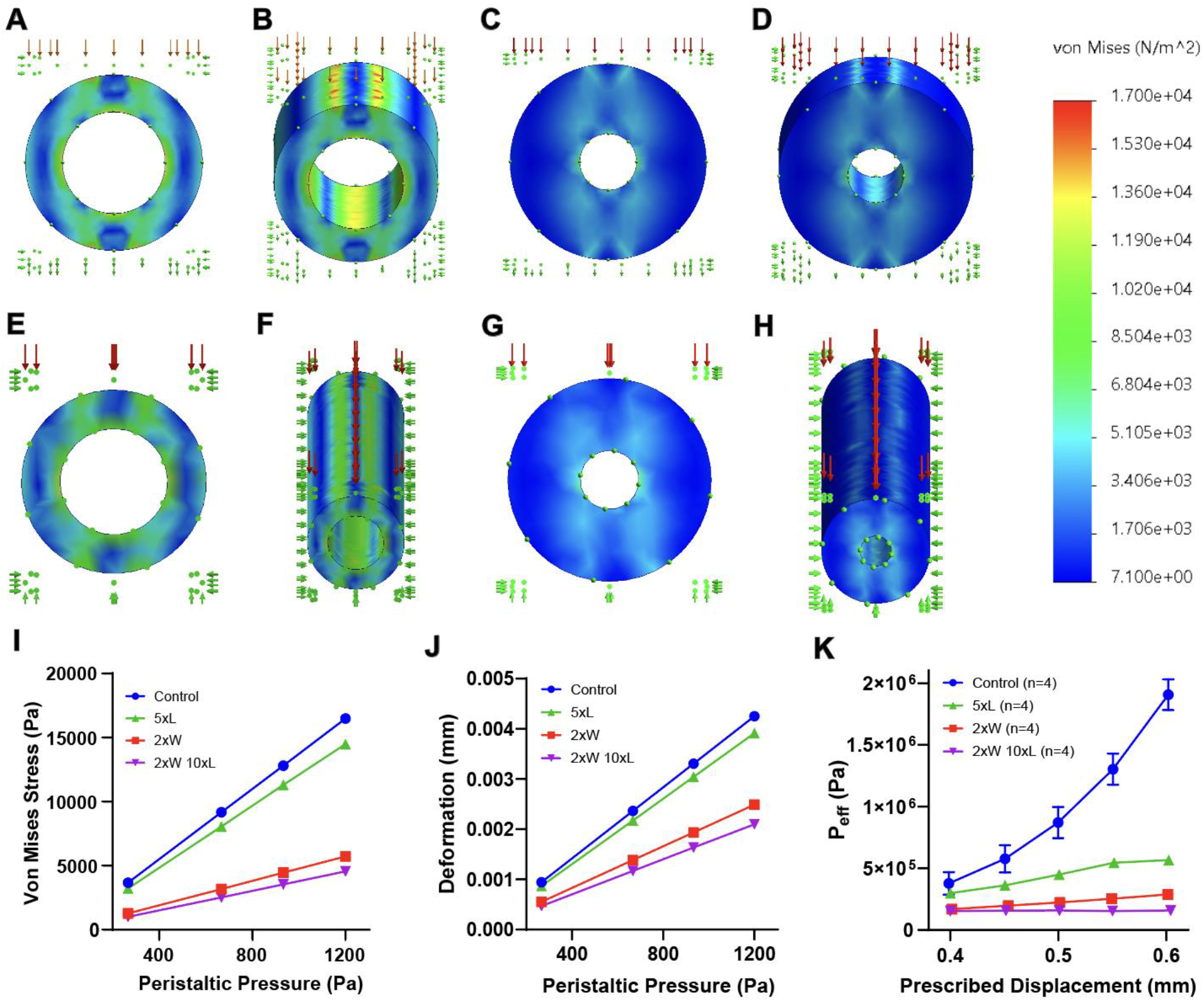
Computational modeling of geometry dependent stress and deformation. (A–H) Finite element simulations showing von Mises stress distributions for control, 5xL, 2xW, and 2xW 10xL marker geometries under physiologically relevant peristaltic pressures. (I) Peak deformation as a function of applied pressure, demonstrating reduced stress and deformation with increasing marker width and length. (J) Peak deformation as a function of applied peristaltic pressure for each marker geometry obtained from finite element simulations. Deformation increases with pressure for all designs, while width modified markers exhibit reduced deformation relative to length only and control geometries. (K) Effective pressure derived from Instron compression testing as a function of prescribed displacement for each marker geometry. Compressive force was converted to effective pressure using an estimated contact area, and the resulting pressure-displacement trends follow the same geometric ranking observed in the simulation based stress and deformation analyses.

Based on the stress and deformation analyses, we hypothesized that increased marker width reduces von Mises stress and overall deformation under peristaltic loading, resulting in mechanically stiffer structures that are less prone to conforming to intestinal contractions and could therefore exhibit prolonged gastrointestinal residency.

To experimentally validate the trends predicted by the computational simulations, we performed uniaxial compressive load testing on the marker geometries using a universal mechanical testing system. Markers were subjected to displacement-controlled compression at a constant rate of 6mm/min, and the resulting compressive force was recorded as a function of prescribed displacement (Figure 2L). To enable comparison with the simulation results, which report stress and pressure based metrics, the measured force was converted to an effective pressure experienced by the marker using:

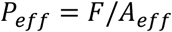

Where F is the measured compressive force and *A*_*eff*_ is the estimated contact area, approximated as the product of the marker length and outer diameter (*A*_*eff*_ = *L*_*marker*_* *d*_*marker*_).

Across the tested displacement range, the experimentally derived effective pressures followed the same geometric ranking observed in the finite element simulations of peak von Mises stress and deformation, with the ordering: control > 5xL > 2xW > 2xW 10xL. While the absolute magnitudes differ due to the simplified loading and contact assumptions inherent to the experimental setup, the consistency in relative response across geometries supports the simulation-derived conclusion that increased marker width reduces mechanical loading under compression, and that combined increases in width and length yield the most robust structures under peristaltic like compression.

Collectively, we hypothesized that marker retention time is governed by the combined effects of marker width, which modulates mechanical deformability under peristaltic loading, and marker length, which influences axial drag and resistance to peristaltic transport, rather than by a single geometric parameter alone.

### 3.2 Radiopacity Comparison Between BaSO_4_-Loaded and Non-Loaded Markers

We next sought to fabricate devices for in vivo testing. Four geometrically distinct marker designs were additively manufactured to assess the influence of size and shape on GI transit (Figure 3A-H). To evaluate the effect of BaSO_4_ incorporation on CT contrast, we compared markers printed with 35% BaSO_4_ concentration to identical markers without BaSO_4_. Overall, microCT imaging showed that BaSO_4_-loaded markers exhibited high radiodensity, appearing as high contrast objects on CT scans across all geometries (Figure 3J, 3L, 3N, and 3P). In contrast, unloaded markers generated minimal signal and were often indistinguishable from the surrounding material under identical scan settings (Figure 3I, 3K, 3M, and 3O).

**Figure 3.**
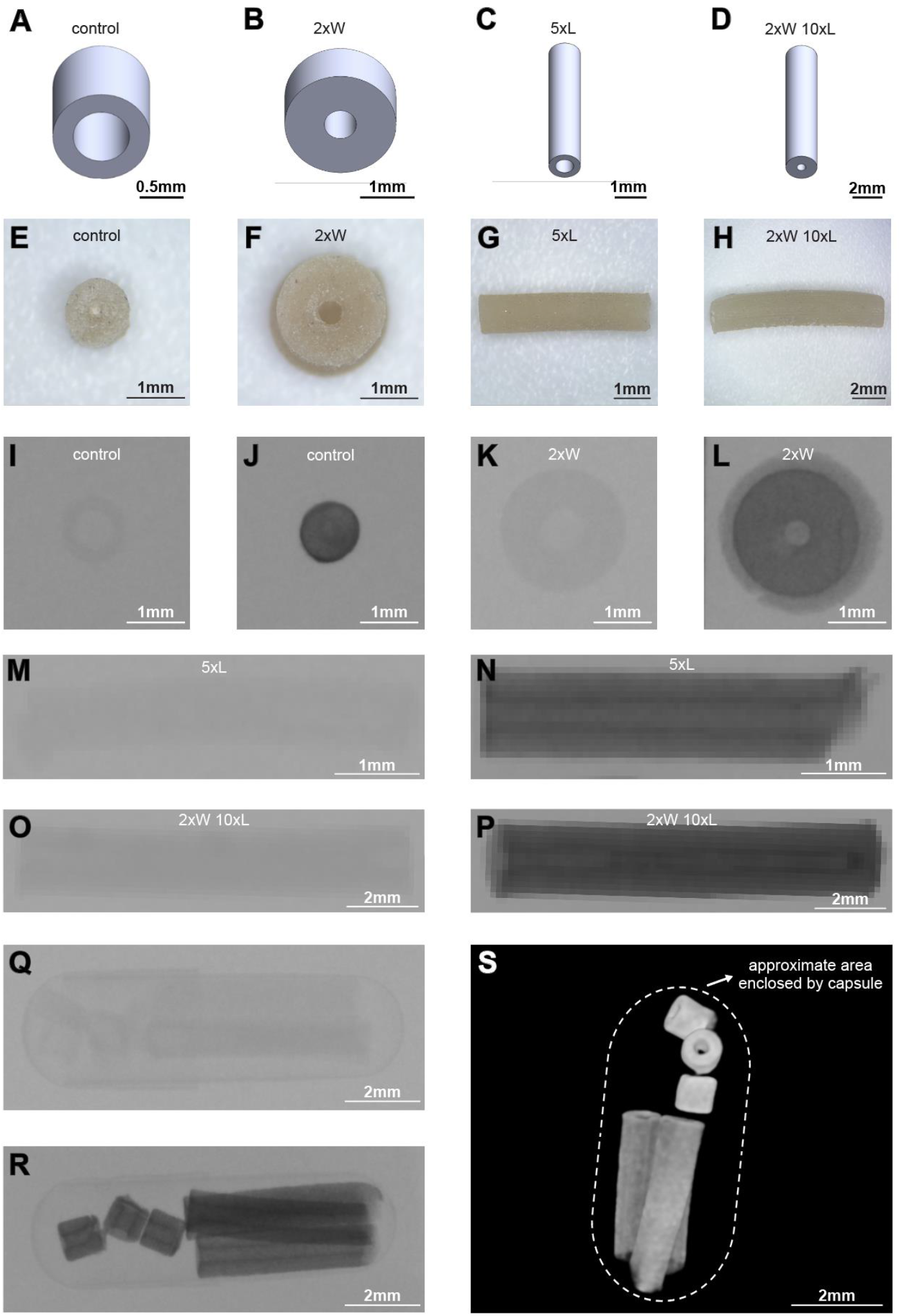
Design, fabrication, and radiopacity of BaSO_4_ loaded 3D printed markers. (A–D) CAD renderings of the control, 2xW, 5xL, and 2xW 10xL marker geometries. (E–H) Optical microscopy images of the corresponding fabricated markers. (I–P) Representative microCT images comparing non BaSO_4_ loaded and 35% BaSO_4_ loaded markers across geometries, demonstrating enhanced radiopacity with BaSO_4_ incorporation. (Q–R) CT images of markers packaged within a gelatin capsule, showing distinguishable geometries prior to administration. (S) Three dimensional microCT reconstruction highlighting the approximate volume enclosed by the capsule and the packed markers.

We packed three control and three 5xL markers into size 9 capsules for in vivo administration to rats (Figure 3Q-S). These images confirmed that multiple BaSO_4_-loaded marker geometries remain clearly visible and readily distinguishable within a single capsule. This further demonstrates the enhanced contrast provided by the incorporation of BaSO_4_ and that loaded markers support reliable identification during multi-marker delivery.

### 3.3 In Vivo MicroCT Tracking from Ingestion to Excretion

We evaluated our hypotheses in vivo using the fabricated devices. We administered BaSO_4_-loaded markers via oral gavage and visualized within the GI tract in vivo at all imaging time points of 1h, 7-9h, 24h, and 48h post-ingestion (Figure 4A-B). At the initial 1h timepoints, markers typically appeared as a compact cluster in the stomach. By 7-9h, the markers began to disperse as they progressed through the gastrointestinal tract. At 24-48h, the distribution became more dispersed, reflecting continued forward movement. Most markers were cleared by 48h, with full elimination generally occurring by 72h. Complete clearance was confirmed through post-excretion microCT scans (Figure 4B), where recovered markers remained distinctly visible in CT. These results demonstrate that BaSO_4_ doped, SLA printed markers can be reliably and noninvasively tracked throughout their entire transit, from ingestion to full excretion, using serial CT imaging.

**Figure 4.**
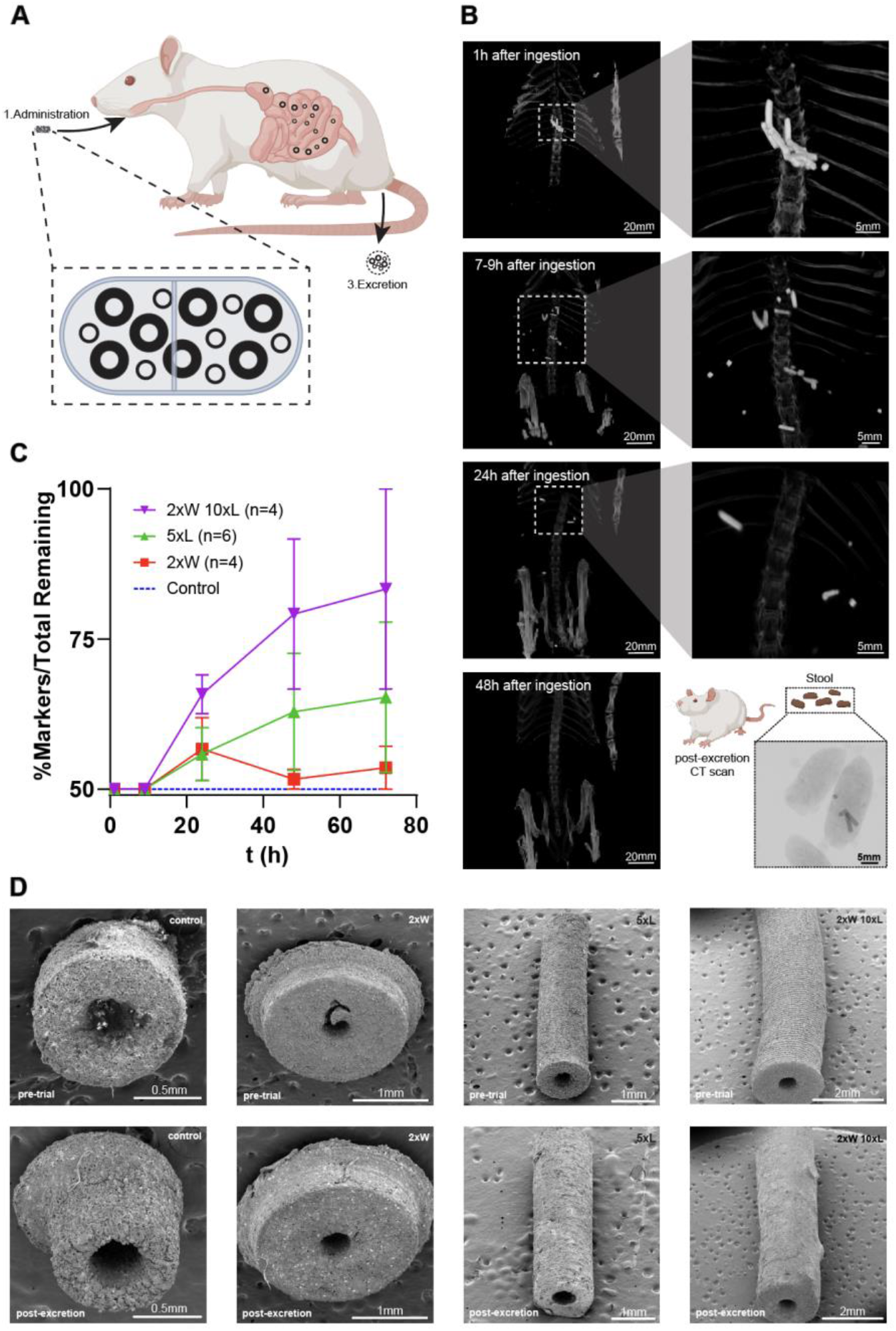
In-vivo MicroCT tracking and geometry dependent gastrointestinal retention. (A) Schematic illustration of capsule loading and oral gavage in the rat model.(B)Representative three-dimensional microCT reconstructions at 1 h, 7–9 h, 24 h, and 48 h post-ingestion, showing progressive marker transit through the gastrointestinal tract and post-excretion confirmation. (C) Quantification of in-vivo retention expressed as the percentage of experimental markers relative to the total number of markers remaining over time, revealing geometry-dependent differences in gastrointestinal residency. (D) Scanning electron microscopy images of markers before administration and after excretion, demonstrating preservation of structural integrity across all geometries.

Across each trial, animal received equal numbers of control and experimental markers in a single capsule. We utilized the relative excretion time for control and experimental geometries to establish a relative retention metric. The retention metric reports the percentage of experimental markers remaining out of the total markers remaining at each time point. If both marker types cleared at the same rate, this value would remain near 50% (control) throughout the experiment (Figure 4C). Deviations above 50% therefore indicate that the experimental geometry remained in the GI tract longer than the control in the same animal.

Quantitative differences in in vivo retention emerged after 24h and varied systematically with marker geometry. At 24h post ingestion, the average percentage of experimental markers remaining was 65.8% for the 2xW 10xL group, compared to 55.8% for 5xL and 56.7% for 2xW markers. These differences became more pronounced over time as control markers were excreted. By 48h, relative retention increased to 79.2%, 62.9%, and 51.7% for 2xW 10xL, 5xL, and 2xW markers respectively. At 72h, the longest and thickest geometry exhibited the greatest persistence, with an average 83.3% of remaining markers belonging to the 2xW 10xL group, compared to 65.3% for 5xL and 53.6% for 2xW. Overall, the data demonstrates a geometry dependent increase in gastrointestinal residency, with the ranking: 2xW 10xL > 5xL > 2xW > control, where the longest and thickest geometry shows the greatest residency (Figure 4C).

The in vivo results support our hypotheses from the analytical model and computational simulations. Together, these data show that width and length influence different mechanical aspects of transit. The width modified markers display lower stresses under compression, consistent with lowered deformability, whereas changes in length have a comparatively smaller effect on deformation behavior. Length, however, contributes to axial drag as described by the analytical model introduced earlier. This distinction helps explain the in vivo retention patterns: the 5xL marker remains longer than the 2xW marker despite being more deformable, indicating that deformability alone does not account for the observed hierarchy. Instead, both deformation behavior and axial drag contribute to overall residency, with the 2xW 20xL design exhibiting the greatest retention because it combines lowered deformability from its larger width with higher axial drag arising from its length.

### 3.4 Marker Geometry and Structural Preservation

Lastly, to verify that GI exposure did not damage the printed structures, we compared pre-trial and post-excretion markers using SEM imaging (Figure 4D). All markers retained their cylindrical shape, hollowcore, and surface features, with no measurable swelling, cracking, or degradation. These results confirm that the 35% BaSO_4_ resin formulation maintains structural integrity during the full ingestion to excretion cycle.

## 4. Discussion

Accurate measurement of gastrointestinal transit and motility remains challenging, and existing clinical and preclinical methods each involve important tradeoffs. Clinically, sitz markers are widely regarded as the first-line technique for evaluating general GI dysmotility. Scintigraphy can also assess gastric emptying and regional gastrointestinal transit, as it enables noninvasive, time resolved tracking of a radiolabeled meal through the GI tract and provides quantitative motility metrics (standard gastric emptying and GI transit scintigraphy)^9^. In preclinical studies, however, simpler assays are more commonly employed and are often terminal in nature. For example, red dye based gastric emptying tests involve oral administration of a colored tracer followed by animal sacrifice and volumetric quantification of dye remaining in the stomach at a single endpoint, rather than dynamic assessment of transit (terminal dye based gastric emptying assays)^10^. Other commonly used approaches, such as the colonic bead or fecal pellet propulsion test, are restricted to evaluating distal colonic motility and do not capture whole gastrointestinal transit behavior^11^. While these methods are valuable for addressing specific physiological questions, they are inherently limited by their terminal design, anatomical restriction, or lack of longitudinal resolution. These limitations reflect the need for platforms that enable noninvasive, time resolved tracking of GI transit across the entire gastrointestinal tract in the same animal. Our platform enables longitudinal, noninvasive CT tracking of ingestible markers in live animals from ingestion through complete excretion, allowing direct measurement of time resolved residency profiles and geometry dependent behavior during gastrointestinal transit. This distinction is particularly important for studying motility, where dynamic progression and clearance kinetics, rather than endpoint localization alone, are critical readouts.

Our goal was to establish a customizable, 3D printed radiopaque marker platform for noninvasive tracking of gastrointestinal (GI) transit and to investigate how simple geometric changes influence residency time in vivo. The ability to manufacture radiopaque geometries with additive manufacturing was crucial. Others have used zirconium oxide (ZrO_2_) nanoparticles in UV curable resin, or polycaprolactone (PCL) with barium sulfate (BaSO_4_)^12,13^, to create biodegradable implants and stents that can be monitored radiographically as well as personalized X-ray shields for targeted tumor irradiation^13,14^.

We combine mathematical modeling, finite element simulations, and in vivo experiments with longitudinal CT imaging in live rats, we provide a unified examination of how marker design shapes transit behavior. These results highlight the feasibility of using BaSO_4_ doped SLA printing for in vivo motility research while also revealing the distinct roles that marker geometry play in determining GI residency.

A simplified mathematical model suggested that axial drag increases proportionally with marker length, while diameter becomes physiologically capped by the maximum distension of the lumen. Under this constraint, diameter contributes relatively little to drag once the marker approaches the near occlusive range, leading to the prediction that length should be the dominant geometric factor influencing resistive forces during transit. Width, in contrast, was expected to influence deformability rather than drag.

The simulation results supported this distinction. Increasing width substantially reduced the stresses experienced during axial compression, demonstrating that width primarily affects deformation behavior, whereas increasing length produced smaller changes in deformation. The in vivo retention patterns aligned with this division of roles: the longest and thickest marker (2xW 10xL) remained in the GI tract for the greatest duration, followed by the length-increased marker (5xL), then the width increased marker (2xW), with the control clearing the fastest. The fact that the 5xL marker outlasted the less deformable 2xW marker shows that deformability alone does not govern retention. Rather, the interplay between deformation behavior and axial drag determines overall residency, and the 2xW 10xL design stays longest because it combines both lowered deformability from its larger width, and increased axial drag from its greater length.

Overall, the results demonstrate that even modest changes in geometry can shift transit behavior in predictable ways, with length and width shaping residency through distinct mechanical pathways. This tunability opens opportunities for creating application specific markers or ingestible devices whose transit profiles can be engineered for either rapid clearance or prolonged retention. The ability to pair customized 3D printed geometries with radiopaque contrast further enables in vivo validation of these design principles in real time.

Future work could explore incorporating a wide array of geometric features and topographies into ingestible markers. Understanding the transit of markers would benefit from more advanced analytic, computational, and probabilistic models. For example, our axial compression model represents a simplified approximation of peristaltic loading; incorporating circumferential and asymmetric contraction patterns could refine the mechanical predictions. Our current model also assumed a parallel alignment of markers with the intestinal lumen. While this may hold for wide markers whose diameter is similar to the luminal diameter, smaller markers are likely constantly re-oriented by contractions and the presence of other ingesta in the gut. Utilizing alternative imaging modalities (e.g. ultrasonic, optical) may also allow for real-time tracking of markers, without sequential radiographs.

This work demonstrates a flexible platform for creating customized radiopaque markers and provides a mechanistic foundation for understanding how simple geometric features govern GI transit. By coupling computational modeling with in vivo imaging, our approach offers a design driven framework for engineering the next generation of ingestible devices and transit modulated delivery systems.

## Acknowledgements

We acknowledge New York University Abu Dhabi Core Technology Platforms for equipment and technical assistance from Dr. Oraib Al-Ketan and Dr. James Weston.

